# The C-terminus of the KREH1 helicase is important for RNA binding and association with mitochondrial RNA editing complexes in *T. brucei*

**DOI:** 10.64898/2026.09.29.753996

**Authors:** Rutuja Yelmar, Laura Tengo, Frank Stein, Eva Kowalinski

## Abstract

Mitochondrial pre-mRNA editing in kinetoplastids, a clade that includes human-infective parasitic protists such as *Trypanosoma* and *Leishmania*, is required to produce translatable, functional gene products from otherwise nonfunctional mitochondrial precursor transcripts of essential respiratory chain complex subunits. RNA editing is a template-guided process in which cognate guide RNAs (gRNAs) anneal to specific sites at the pre-mRNA and direct the complementary insertion and/or deletion of uridines. Large multi-modular complexes - like the <u>R</u>NA-<u>E</u>diting <u>S</u>ubstrate-binding <u>C</u>omplex (RESC) and the <u>R</u>NA-<u>E</u>diting <u>C</u>atalytic <u>C</u>omplexes (RECC) - organize and catalyze RNA editing, respectively. Within these assemblies, <u>K</u>inetoplast <u>R</u>NA <u>E</u>diting <u>H</u>elicases (KREHs) play an important role in ribonucleoprotein complex remodeling, particularly in pan-editing, when transcripts are extensively modified throughout their length by multiple gRNAs. However, the mechanisms by which KREH RNA helicases facilitate pan-editing, and their specific roles in this process, remain unclear. Here, we define the molecular properties of the *Trypanosoma brucei* DEAD-box RNA editing helicase KREH1. Cellular co-proteome analyses reveal that KREH1 interacts with subunits of the RESC complex and biochemical experiments show that KREH1 is an RNA-dependent ATPase that unwinds double-stranded RNA substrates. We show that the flexible C-terminus of KREH1 is important for RNA binding and, consequently, for enzymatic activity *in vitro* and interactions with the RESC complex *in vivo*. Overall, our experiments refine the role of KREH1 in the pan-editing process and establish its utility as a molecular tool for investigating higher-order RNA editing complexes in the cellular context.

## Introduction

Kinetoplastids, a clade of parasitic protists including *Trypanosoma* and *Leishmania* genera, cause vector-borne diseases in humans and animals (Stuart et al. 2008). The kinetoplast, the name-giving feature of this clade, is a tightly interwoven network of mitochondrial circular DNAs (kinetoplast DNA, kDNA), attached to the basal body of the flagellum in direct proximity to the flagellar pocket (Ogbadoyi et al. 2003). The kDNA consists of 20 to 30 ‘maxicircles’, approximately 20–40 kilobases (kb) in size, which encode 9S and 12S rRNA, two ribosomal proteins, and subunits of mitochondrial respiratory chain complexes (reviewed in Lin et al. 2015). However, most maxicircle gene sequences are incomplete, and the primary transcripts do not encode functional protein. Instead, these cryptic precursors, also referred to as pre-edited mRNAs, must undergo additional processing by insertion and/or deletion of uridine bases, through a process known as kinetoplastid mitochondrial mRNA editing, to ultimately encode intact protein sequences (Feagin et al. 1988; Shaw et al. 1988; Benne et al. 1986). The editing is templated by another RNA species, small 50–60 nucleotide (nt) long gRNAs encoded by thousands of kDNA ‘minicircles’, which are approximately 1 kb in size (Lukeš et al. 2002). For gRNA biogenesis, a sense-antisense duplex is generated via bidirectional transcription, the 3′ end of the sense strand is polyuridylated by the ‘mitochondrial 3′ processome’ (MPsome), and the antisense strand is degraded (Suematsu et al. 2016; Aphasizhev et al. 2016). The mature gRNA carries a 5′-triphosphate and a 10–12 nt anchor region matching its target pre-edited mRNA through reverse-complementary base-pairing. In its central region, the gRNA provides the corrective template sequence for uridine insertion/deletion editing, while its 3′ end is marked by a poly(U) tail (reviewed in Aphasizheva et al. 2020; Stuart et al. 2005). As a prerequisite for the editing reaction, pre-mRNA and gRNAs anneal, and mismatches form bulge structures that attract the <u>R</u>NA <u>E</u>diting <u>C</u>atalytic <u>C</u>omplexes (RECC). The RECC complexes share common scaffolding subunits and different endo- or exonucleases, terminal uridylyl transferase (TUTase), and ligases to cleave the pre-mRNA, insert and/or remove uridines, and re-ligate it (reviewed in Aphasizheva et al. 2020; Liu et al. 2026). Modules of the large <u>R</u>NA <u>E</u>diting <u>S</u>ubstrate Binding <u>C</u>omplex (RESC) form essential scaffolds that coordinate mRNA, gRNAs, and RECC; the approximately 20 subunits of RESC do not possess catalytic activity (Aphasizheva et al. 2014; Dolce et al. 2023; Dubey et al. 2021; Liu et al. 2023; Panigrahi et al. 2007; Weng et al. 2008).

Depending on the extent of editing, mitochondrial mRNAs are classified as ‘never edited’ (CO- Cytochrome oxidase, COI; ND-NADH dehydrogenase subunit, ND1, ND4, ND5; MURF-maxicircle unidentified reading frame, MURF1, MURF5), ‘minimally edited’ (COII; Cyb-Cytochrome b; MURF2), or ‘pan-edited’, when extensive editing is required across their entire sequence (ATPase subunit 6 (A6); COIII; C-rich reading frame 3 and 4 (CR3, CR4); Ribosomal protein 12 (RPS12); ND3, ND7, ND8, ND9) (reviewed in Hajduk and Ochsenreiter 2010). By definition, a single gRNA directs modification of an editing ‘block’ leading to editing of one or more insertion and/or deletion sites (Blum et al. 1990). In some cases, restoring the start codon serves as a marker for the completion of editing (Shaw et al. 1988); alternative editing allows for expression of protein variants, enhancing proteome diversity (Gerasimov et al. 2022; Ochsenreiter and Hajduk 2006). Pan-editing involves multiple, sometimes overlapping gRNAs when extensive editing spans multiple blocks and typically proceeds from the 3′ to the 5′ end of the mRNA (Maslov and Simpson 1992). RNA editing relies on the motor activities of auxiliary <u>K</u>inetoplast <u>R</u>NA <u>E</u>diting <u>H</u>elicases (KREHs) (reviewed in Aphasizheva et al. 2020; Stuart et al. 2005; Cruz-Reyes et al. 2016; McDermott et al. 2026).

RNA helicases fulfill diverse functions in cells, with ATP-dependent activities ranging from RNA binding to unwinding or remodeling of RNA structures (reviewed in Pyle 2008). In eukaryotes, mitochondrial RNA helicases play pivotal roles. For example, Suv3p, a DExH-box RNA/DNA helicase from *Saccharomyces cerevisiae*, is involved in the degradation of excised group I intron RNAs (Margossian et al. 1996; Turk and Caprara 2010), while the DEAD-box RNA chaperone, CYT19 from *Neurospora crassa,* facilitates group I intron splicing (Busa et al. 2017; Mohr et al. 2002). In the mitochondria of kinetoplastids, the DEAD-box helicase KREH1 (formerly Hel61 or REH1) facilitates RNA editing, potentially by resolving mRNA structures, facilitating the removal of RNA-binding proteins, or displacing gRNAs from the gRNA:mRNA duplex upon completion of editing. The involvement of KREH1 in kinetoplastid RNA editing was first demonstrated through the helicase activity in mitochondrial extracts (Corell et al. 1996; Missel et al. 1997; Missel and Göringer 1994). Knockout of KREH1 in the procyclic form *T. brucei* leads to reduced levels of edited mRNAs, whereas the abundance of never-edited mitochondrial transcripts and nuclear-encoded mRNAs remains unchanged, suggesting that the protein is nonessential and/or that functionally redundant enzymes exist (Dubey et al. 2023; Missel et al. 1997). Co-localization analysis and *in vivo* co-immunoprecipitation analysis of complexes identified the association of KREH1 with mitochondrial RNA editing complexes; several lines of evidence suggest that this interaction is transient and/or dependent on RNA (Dubey et al. 2023; Hashimi et al. 2008; Panigrahi et al. 2003, 2006; Li et al. 2011). More detailed studies have identified KREH1 as a factor that facilitates editing progression, particularly in cases of overlapping guide RNAs (gRNAs), pointing towards a role in post-editing duplex resolution. Repression of KREH1 expression led to increased abundance of the gRNA:edited mRNA duplex of the first editing block in A6 mRNA (Li et al. 2011). High-throughput sequencing of catalytically inactive KREH1 cell lines suggests a role for KREH1 in editing initiation (Dubey et al. 2023). Recombinant *Leishmania major* KREH1 is an active ATP-dependent DEAD-box helicase unwinding RNA duplexes (Li et al. 2011). To date, the precise substrate preferences and molecular basis of the interaction between *T. brucei* KREH1 and RNA remain uncharacterized.

In this study, we elucidate the molecular function of the KREH1 RNA helicase from *T. brucei*, focusing on its structural features, substrate preferences, and interactions with subunits of the RNA editing machinery. We identify a role for the KREH1 flexible C-terminal region in its function as RNA helicase: the C-terminus is necessary for RNA binding *in vitro* and for its association with the editing machinery *in vivo*. We systematically probe *T. brucei* KREH1 activity across various substrates and demonstrate that the enzyme functions as an RNA-dependent ATPase with a preference for double-stranded (ds)RNAs, carrying mismatches and 3′ overhangs. In addition, using molecular and cellular experiments, we establish the use of ATP analog inhibitors to stabilize KREH1 interactions with RNA substrates and proteins of the editing machinery. These findings enhance our understanding of editing-associated RNA helicases and provide a tool for tethering RNA editing complexes *in vivo,* thereby supporting future *in situ* structural studies.

## Materials and methods

### KREH1 structure prediction

The AlphaFold2 model of KREH1 (Tb927.11.8870) was generated with AlphaFold2 (v2.0.0) on Google Colab (Jumper et al. 2021; Mirdita et al. 2022). Models were visualized with ChimeraX (daily builds, UCSF) (Pettersen et al. 2021).

### Construct design and cloning

All plasmids and oligonucleotides used in this study are detailed in Supplementary Tables S1 and S2, respectively. For co-proteome determination, KREH1 constructs were amplified from *T. brucei* AnTat 1.1 genomic DNA and cloned into the pLEW82v4 vector for genomic integration into the rRNA spacer locus (Wirtz et al. 1999). Constructs were fused at the N-terminus to the native KREH1 mitochondrial targeting sequence (MTS; MRALRCVRRGVYRQSVRLCYFMSLECSLR), predicted using MitoFates (Fukasawa et al. 2015), followed in-frame by an enhanced green fluorescent protein (eGFP)– 3C protease cleavage site (3C)–streptavidin-binding peptide (SBP) tag. For recombinant expression in *E. coli*, KREH1 constructs were cloned into the pEC vector (Elena Conti, Max Planck Institute of Biochemistry, Germany) with an 8xHis-3C protease amino-terminal tag. ATPase-deficient mutants (E269Q) were generated with site-directed mutagenesis PCR.

### *Trypanosoma brucei* culture and cell line generation

*T. brucei SmOx* cells (Poon et al. 2012) were cultured as procyclic form in SDM-79 medium supplemented with 10% (v/v) heat-inactivated fetal bovine serum, 20 µg/ml penicillin-streptomycin, 0.4% (v/v) glycerol, and 30 µg/ml hemin at 27 ℃. Cells were maintained at a density of 5 × 10^6^ cells/ml. Before transfections and other experiments, cells were passaged at least two times after thawing. All cell lines generated are indicated in Supplementary Table S3. For the generation of overexpression cell lines, the pLEW82v4 vector was linearized with NotI (NEB, #R3189S) and purified by ethanol precipitation (mixing 10 µl of 3 M sodium acetate and 300 µl of 100% EtOH, 1h incubation at -80 ℃, washing twice with 70% EtOH). Approximately 10 µg of DNA was transfected into 2 × 10^7^ cells in 500 µl of Cytomix buffer (25 mM HEPES pH 7.6, 120 mM KCl, 0.15 mM CaCl_2_, 10 mM K_2_HPO_4_/KH_2_PO_4_ pH 7.6, 2 mM EDTA, 6 mM glucose, 5 mM MgCl_2_, sterile filtered) by delivering a pulse at 800 V, 50 µF, and 100 Ω using a BTX Gemini SC2 electroporation instrument. Cells were transferred to 5 ml of antibiotic-free conditioned medium and incubated for 24 h before the addition of zeocin (InvivoGen) to a final concentration of 5 µg/ml. After another 7 days, the zeocin concentration was reduced to 2.5 µg/ml. Recovery of positive clones to a doubling time of ∼12 h was achieved after approximately 3 weeks. KREH1 expression was induced by the addition of tetracycline to a final concentration of 1 µg/ml for 24 h.

### Immunoprecipitation

For each sample, 1 × 10^9^ cells were harvested by centrifugation (1,500 × g, 20 min, 4 ℃), washed, and resuspended in 10 ml of immunoprecipitation (IP) buffer (20 mM HEPES pH 7.5, 200 mM NaCl, 10 mM MgCl_2_), freshly supplemented with a Complete protease inhibitor tablet (Roche) and 1 mM DTT. When indicated, 2 mM ADP:BeF_x_ (Sigma) was added in this step (ADP and BeF_x_ complex were mixed in a 1:1 molar ratio; BeF_x_ was obtained by mixing BeSO_4_ and KF in a 1:2 molar ratio). Cells were lysed by sonication (5 s on, 15 s off, for 4 min at 30% amplitude; Vibra-cells Sonics, on ice). The lysate was clarified by centrifugation at 12,096 × g for 60 min at 4 ℃, and the supernatant was divided into three technical replicates, each containing approximately 90 mg of total protein.

Each replicate was incubated with 100 µl of eGFP-nanobody resin (based on CNBr-activated Sepharose 4B resin) for 1 h at 4 ℃ in an orbital shaker; the resin had been pre-equilibrated in IP buffer. The resin was washed twice (900 × g for 1 min at 4 ℃) with 500 µl of IP buffer. For elution, the resin was mixed with 140 µl of IP buffer containing 4 µM of 3C protease (EMBL PEPCF) for 1 h at 4 ℃. 50 µl of the eluate was subjected to TMT mass spectrometry.

### Western blot

The cell lysate from KREH1 overexpression cell lines were generated for immunoprecipitation experiment using the protocol. The total cell lysate (T) and soluble clarified cell lysate (S) were separated using a 12% SDS-PAGE gel with pre-stained Page ruler protein marker (Thermo Fisher Scientific), positive control -MS2-GFP (36 kDa) and negative control untransfected SmOx cell lysate. Blotting membrane PVDF (polyvinylidene difluoride) was activated for 10 min in 100% ethanol. Activated membrane, SDS-PAGE gel, and Whatman filter paper were pre-equilibrated with transfer buffer (190 mM Glycine, 20 mM Tris-Base, 20% Ethanol) for 10 min. The transfer cassette was assembled and blotted at 80 V for 90 min with Bio-Rad Mini-Protean Tetra Vertical Electrophoresis Cell on ice. After blotting, the membrane was blocked with 5% milk prepared in 1× PBST overnight at 4 ℃. A primary mouse anti-GFP antibody (Santa Cruz, sc-9996) was diluted in 1× PBST (0.1% Tween20) with 1% milk (w/v) at a concentration of 1:500 and incubated for 1 h at room temperature, followed by three washes with 20 ml of 1× PBST for 10 min each. The secondary anti-mouse antibody (Cell Signaling 7076) was then prepared in 1× PBST-1% milk (w/v) and applied at a dilution of 1:10000 for 1 h at room temperature, followed by another set of three washes with 20 ml of 1× PBST for 10 min each. The blot was developed using 1 ml of SuperSignal^TM^ West Femto Maximum Sensitivity Substrate (Thermo Scientific^TM^) and visualized on a ChemiDoc (Bio-Rad) with a chemiluminescence filter at various exposure times.

### Microscopy

1 ml of cells at a density of 5 × 10^6^ cells/ml was centrifuged at 1,400 × g for 3 min at 4 ℃. The cells were resuspended in 415 µl of serum-free SDM-79 medium, and 75 µl of 16% (w/v) paraformaldehyde was added. The mixture was then incubated for 16 h at 4 ℃ in the dark. Cells were centrifuged at 1,400 × g for 3 min at 4 ℃ and washed with 1 ml of phosphate-buffered saline (PBS) before resuspension in 20 µl of PBS. 2 µl of cells and 2 µl of 1:1000 NucBlue stain (Thermo Scientific) were placed on a microscope slide, and a coverslip was applied before imaging with a 60× oil immersion objective on Evos imager (Thermo Scientific) via GFP and DAPI channels. The resulting images were processed with Fiji (Schindelin et al. 2012).

### Recombinant protein production and purification

KREH1 recombinant constructs were transformed and expressed in *E. coli* strain Rosetta 2 (DE3) cells (Novagen/Merck) cultivated in Terrific Broth (TB) medium supplemented with ampicillin or kanamycin to a final concentration of 100 µg/ml or 50 µg/ml, respectively and chloramphenicol up to 34 µg/ml. Protein expression was induced at OD_600_ 0.8 with 0.2 mM isopropyl-1-thio-β-D-galactopyranoside (IPTG) for 18 h at 18 ℃. The cells were harvested via centrifugation at 4,000 × g for 20 min at 4 ℃ and stored at -80 ℃. Pellets were resuspended in lysis buffer (20 mM Tris-HCl pH 7.5, 200 mM NaCl, 20 mM imidazole, and 2% glycerol) freshly supplemented with 0.01% (v/v) beta-mercaptoethanol (≥ 99%), 17.4 µg/ml PMSF, 50 µg/ml lysozyme, 4 µg/ml DNase I, and 40 µg/ml RNase A. Cells were lysed by sonication (5 s on, 15 s off, for 4 min at 40% amplitude, on ice). The lysate was clarified by centrifugation at 25,100 × g for 45 min at 4 ℃ and then applied to a 5 ml nickel-nitrilotriacetic acid (Ni-NTA) HisTrap HP affinity chromatography column (Cytiva), which had been pre-equilibrated in lysis buffer. The resin was washed with 10 ml of low salt buffer (20 mM Tris-HCl pH 7.5, 200 mM NaCl, 50 mM imidazole), followed by washing with 10 ml of high salt buffer (20 mM Tris-HCl pH 7.5, 1000 mM KCl, 50 mM imidazole), and with 40 ml of low salt buffer. The protein was eluted in 10 ml elution buffer (20 mM Tris-HCl pH 7.5, 200 mM NaCl, 600 mM imidazole) and applied to a 5 ml HiTrap Heparin HP affinity chromatography column (Cytiva) pre-equilibrated with low salt buffer; then eluted with a linear gradient to 1 M NaCl in the same buffer. The pooled eluate fractions were treated with 3C protease in a 1:50 (w/w) ratio protease: protein. The protein was concentrated using a 30 kDa MWCO Amicon filter (Merck Millipore) and aliquots were flash frozen in liquid nitrogen and stored at -80 ℃. The final size-exclusion chromatography was performed on a Superdex 200 Increase 3.2/300 GL column (Cytiva) in SEC buffer (20 mM HEPES pH 7.5, 200 mM NaCl, 2 mM MgCl_2_, and 1 mM DTT). The purification protocol remained consistent for all constructs, except for KREH1ΔN99, which utilized 250 mM NaCl in all buffers. Of note, recombinant KREH1 was aggregation-prone, sensitive to buffer conditions below 100 mM NaCl, and could not be concentrated beyond 3 mg/ml, thereby limiting the concentration and buffer ranges in subsequent assays.

### RNA duplex preparation

RNA duplexes for the fluorescence polarization and ATPase or unwinding assays were prepared in ultra-pure water or buffer (20 mM HEPES at pH 7.5 and 100 mM NaCl), respectively. Fluorescein amidite (FAM)-labeled and unlabeled synthetic RNA oligomers (biomers.net) were mixed in a 1:1.1 molar ratio in water or buffer, as indicated. The mixture was heated to 95 ℃ for 5 min, then allowed to cool slowly by turning off the heat and allowing the heating block to return to room temperature for approximately 2.5 h. The sequences of all RNA oligos used for biochemical assays are listed in Supplementary Table S4.

### Fluorescence polarization

Serial dilutions of purified ATPase-deficient KREH1 truncations were prepared in protein buffer (20 mM HEPES pH 7.5, 2 mM MgCl_2_, and 250 mM or 200 mM NaCl, depending on truncation solubility). RNA (44 nM) and AMP-PNP (2 mM) were used. The final NaCl concentration was adjusted to 100 mM by adding dilution buffer (20 mM HEPES, pH 7.5, 2 mM MgCl_2_), as KREH1 was prone to precipitate in the absence of RNA at low salt concentrations. Final protein concentrations ranged from 8.7 µM to 2 nM. Polarization measurements were conducted after 5 min of incubation on ice using a Clariostar plate reader (BMG Labtech) equipped with a fluorescein filter. The polarization values from three technical replicates were plotted using GraphPad Prism and analyzed through non-linear regression (total single-site binding). For statistical analysis, the Area Under the Curve (AUC) was calculated for each replicate. Further, the mean AUCs of triplicates were compared using one-way ANOVA with Tukey’s multiple comparison test to determine statistically significant differences in binding of different KREH1 truncations to dsRNA using GraphPad Prism.

### RNA unwinding assay

For the unwinding assays, a mixture of 150 nM protein, 2 mM ATP, and 500 nM of unlabeled competition DNA oligo (5′ CTATAACTCCAATG 3′) was prepared in reaction buffer (20 mM HEPES pH 7.5, 100 mM NaCl, 2 mM MgCl_2_) and incubated at 30 ℃ for 10 min. The reaction was started by adding 250 nM of RNA substrate. At the indicated time points, aliquots of 10 µl were quenched by mixing with 10 µl of quenching buffer (150 mM NaAc, 10 mM EDTA, 0.025% (w/v) SDS, 25% (v/v) glycerol, 0.25% (w/v) Orange G). Controls were prepared in the same manner but excluded protein or ATP, respectively. Samples were analyzed on 20% native-PAGE (1 ml Tris-acetate EDTA pH 8.3 (10×), 5 ml acrylamide/bis-acrylamide (40%), 250 µl APS (100 mg/ml), 25 µl TEMED (≥ 99%)) and the fluorescence was detected via a ChemiDoc imager (Bio-Rad).

### ATPase assay

The ATP hydrolysis assay was prepared in triplicate in 100 µl reaction volume with the EnzChek Phosphate assay kit (Thermo Fisher), containing 150 nM of protein, 0.2 mM of 2-amino-6-mercapto-7-methyl-purine riboside (MESG) substrate, 0.1 U of purine nucleoside phosphorylase (PNP), 2 mM ATP, in the reaction buffer containing 20 mM HEPES pH 7.5, 100 mM NaCl and 2 mM MgCl_2_. The mixture was incubated at 30 ℃ for 10 min. The reaction was initiated by the addition of 250 nM of the dsRNA substrate, and the release of inorganic phosphate (P_i_) was quantified by measuring absorbance at 360 nm with a Clariostar plate reader (BMG Labtech) at 60 s intervals over 60 min. ATPase activity curves were generated in GraphPad Prism, and ATPase rates were calculated from the slope of the absorbance change at 360 nm during the initial 0–10 min from the curve. Mean ATPase rates were compared for statistical significance using one-way ANOVA with Tukey’s multiple comparison test with GraphPad Prism. The kinetic parameters were assessed at varying RNA concentrations while maintaining a constant protein concentration, utilizing Michaelis-Menten equations in GraphPad Prism.

### Tandem Mass Tag Mass Spectrometry (TMT MS)

Protein samples were processed using the SP3 protocol (Hughes et al. 2014) on the KingFisher Apex™ (Thermo Fisher). Proteins were digested with trypsin (1:20 protease:protein) in 50 mM HEPES, 5 mM TCEP and 20 mM chloroacetamide (CAA) for 5 h at 37 ℃. Up to 10 µg peptides were labeled with TMTpro™ 18plex (Li et al. 2021) following manufacturer instructions, quenched with 5% hydroxylamine, pooled, desalted (Oasis® HLB µElution Plate, Waters), and dried. Peptides were fractionated by high-pH reversed-phase HPLC (Agilent 1200; Gemini C18, Phenomenex (Yang et al. 2012), using buffer A pH 10 containing 20 mM ammonium formate and buffer B containing acetonitrile). A 59-min linear gradient (0–35% B) was applied at 0.1 ml/min, followed by ramp to 85% B and re-equilibration. Forty-eight fractions were collected, pooled into six, and dried. LC–MS/MS was performed on an UltiMate 3000 RSLCnano (Thermo Fisher) with a PepMap™ 100 C18 trap (300 µm × 5 mm) and nanoEase™ M/Z HSS T3 analytical column (75 µm × 250 mm, Waters). Peptides were separated at 0.3 µl/min with 0.1% formic acid in water (A) and acetonitrile (B), both containing 3% DMSO. Data were acquired on a Q Exactive™ Plus (Thermo Fisher) with spray voltage 2.2 kV and 275 ℃ capillary temperature. MS1 scans (m/z 375–1200) were collected at 70,000 resolution; MS2 scans at 35,000 resolution using HCD (NCE 30), 0.7 m/z isolation window, 30 s dynamic exclusion, and charge state 2–4 selection. Raw files were converted with ProteoWizard MSConvert (64-bit, zlib, peak picking, top 1000 peaks) and searched using MSFragger (FragPipe 22.1-build02) against the *Trypanosoma brucei* proteome version 68 available in TriTrypDB with contaminants and decoys. Fixed modifications: Carbamidomethyl (C, +57.0215), TMTpro (K, +304.2072); variable: Oxidation (M, +15.9949), Acetyl (protein N-term, +42.0106), TMTpro (peptide N-term, +304.2072). Trypsin specificity (≤2 missed cleavages, peptide length ≥7) and 20 ppm MS1/MS2 tolerances were used. Peptide/protein false discovery rate (FDR) was controlled at 1%. Quantitative analysis was performed in R (ISBN 3-900051-07-0). Contaminants were removed, retaining 3,625 proteins with ≥2 razor peptides. Replicate batch effects were corrected with removeBatchEffect (Ritchie et al. 2015), followed by variance stabilization normalization with normalizeVSN (Huber et al. 2002). Missing values were imputed (KNN, MSnbase) (Gatto and Lilley 2012). Differential expression was assessed using moderated t-tests using the limma package (Ritchie et al. 2015), with imputed values down-weighted (0.01 vs 1). P-values were adjusted for multiple testing with the Benjamini-Hochberg procedure. No missing values were present in this dataset – all 3,625 protein groups were quantified in all 18 TMT channels. Proteins were annotated as hits at a false discovery rate (FDR) ≤ 0.05 and |fold change| ≥ 2, and as candidates at FDR ≤ 0.2 and |fold change| ≥ 1.5. Results were visualized as volcano plots (GraphPad Prism). The experiment was carried out on triplicates from the same cell lysate i.e. technical replicates thus the reported FDRs describe technical reproducibility rather than biological. The mass spectrometry proteomics data have been deposited to the ProteomeXchange Consortium via the PRIDE (Perez-Riverol et al. 2022) partner repository with the dataset identifier PXD067570.

## Results

### The AlphaFold2 model of KREH1 predicts a DEAD-box helicase fold with N- and C-terminal intrinsically disordered regions

To investigate the molecular determinants guiding KREH1 recruitment to specific protein and RNA components of the kinetoplastid mitochondrial mRNA editing machinery, we generated an AlphaFold2 model of KREH1 (Jumper et al. 2021; Mirdita et al. 2022). The KREH1 model reveals a classical DEAD-box helicase core comprising two RecA-like domains with intrinsically disordered N- and C-terminal appendices (Figure 1A-C; Supplementary Figure S1A). A Foldseek search (van Kempen et al. 2024) identified the multifunctional human DEAD-box RNA helicase DDX3X (E = 1.662 × 10^-37^) (Lai et al. 2016; Lee et al. 2008) and the ribosomal RNA processing DEAD-box RNA helicase DbpA from *E. coli* (E = 1.101 × 10^-30^) (Diges and Uhlenbeck 2001; Henn et al. 2010), as the closest structural homologs of KREH1. In the 100-residue-long N-terminal extension, a mitochondrial targeting signal (MTS) is predicted by MitoFates (Fukasawa et al. 2015). The C-terminal tail is shorter, ranging from 27–45 amino acids in *Trypanosoma* and *Leishmania*, respectively, and harbors conserved positively charged residues that may mediate interactions with proteins or RNA (Figure 1C; Supplementary Figure S1B, C).

**Figure 1.**
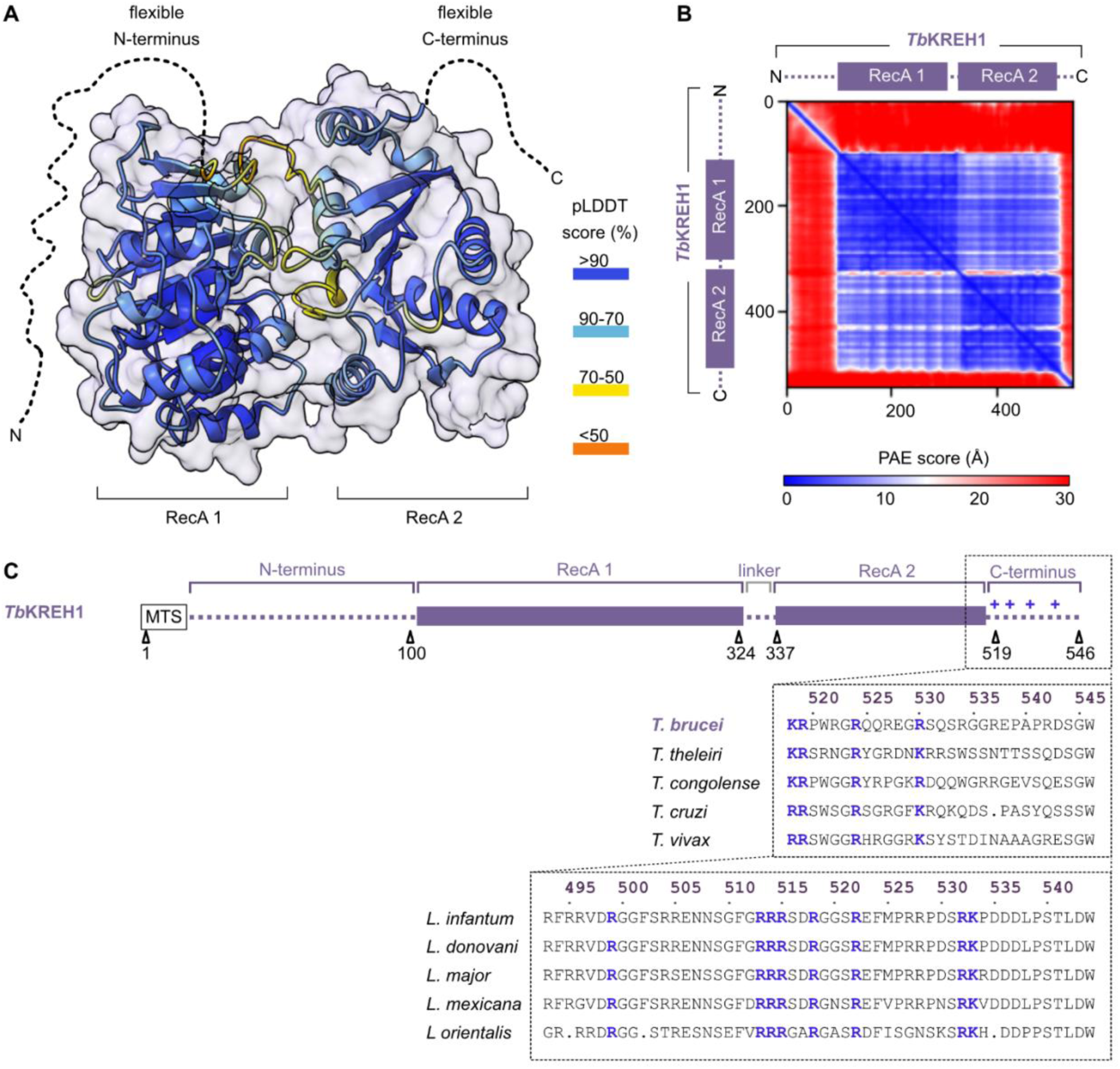
Structural model of KREH1. **(A)** The T. brucei KREH1 AlphaFold2 model in cartoon and surface representation for the RecA core. The cartoon is colored by per-residue confidence (pLDDT, %). The disordered amino- and carboxy-termini are shown as dashed lines (pLDDT < 50%). (**B**) Predicted Aligned Error (PAE) plot indicating pairwise positional uncertainty (blue: low, red: high). (**C**) Annotated linear schematic of T. brucei (Tb) KREH1: the mitochondrial targeting signal peptide (MTS), the helicase core comprising two RecA-like domains, and a flexible carboxy-terminus with positively charged residues (blue) conserved within Trypanosoma species and in Leishmania species; dotted lines represent predicted disordered regions.

### ATPase inhibitor-induced helicase stalling enhances the cellular KREH1 interactome

To assess the cellular interactome of KREH1 and identify interactions that specifically depend on its N- and C-terminal unstructured regions, we tagged KREH1 for inducible expression in *T. brucei* and performed co-immunoprecipitation followed by proteomic analysis. To ensure correct localization of our constructs, we added the original predicted KREH1 mitochondrial targeting sequence (MTS) N-terminal to the affinity tag (a fusion of enhanced green fluorescent protein, 3C protease cleavage peptide, and streptavidin binding peptide, MTS-eGFP-3C-SBP). The cell lines expressed a full-length version - KREH1FL (residues P30-W546) - and two variants with truncated termini, lacking either the N-terminal 99 residues - KREH1ΔN99 (residues D100-W546) - or the C-terminal 27 residues - KREH1ΔC27 (residues P30-R519) (Figure 2A). Fluorescence imaging demonstrates the correct localization of the tagged constructs to the mitochondrion, and western blots indicate the presence of the tagged proteins in the soluble fraction that we used for immunoprecipitation on eGFP nanobody resin (Figure 2B; Supplementary Figure S2). We analyzed the co-purified proteome using tandem mass tag (TMT) multiplex mass spectrometry (MS) for protein identification and comparative enrichment analysis, with untagged cells as a control. Despite successful enrichment of tagged KREH1, the initial enrichment of co-precipitating proteins was weak, with tagged KREH1FL samples nearly indistinguishable from control cells lacking the tagged protein (Figure 2C, Supplementary Table S5). This observation is consistent with prior experiments, which show that KREH1 association with the mRNA-editing complexes was weak, possibly transient, and RNA-dependent (Hashimi et al. 2008; Panigrahi et al. 2003, 2006; Li et al. 2011).

**Figure 2.**
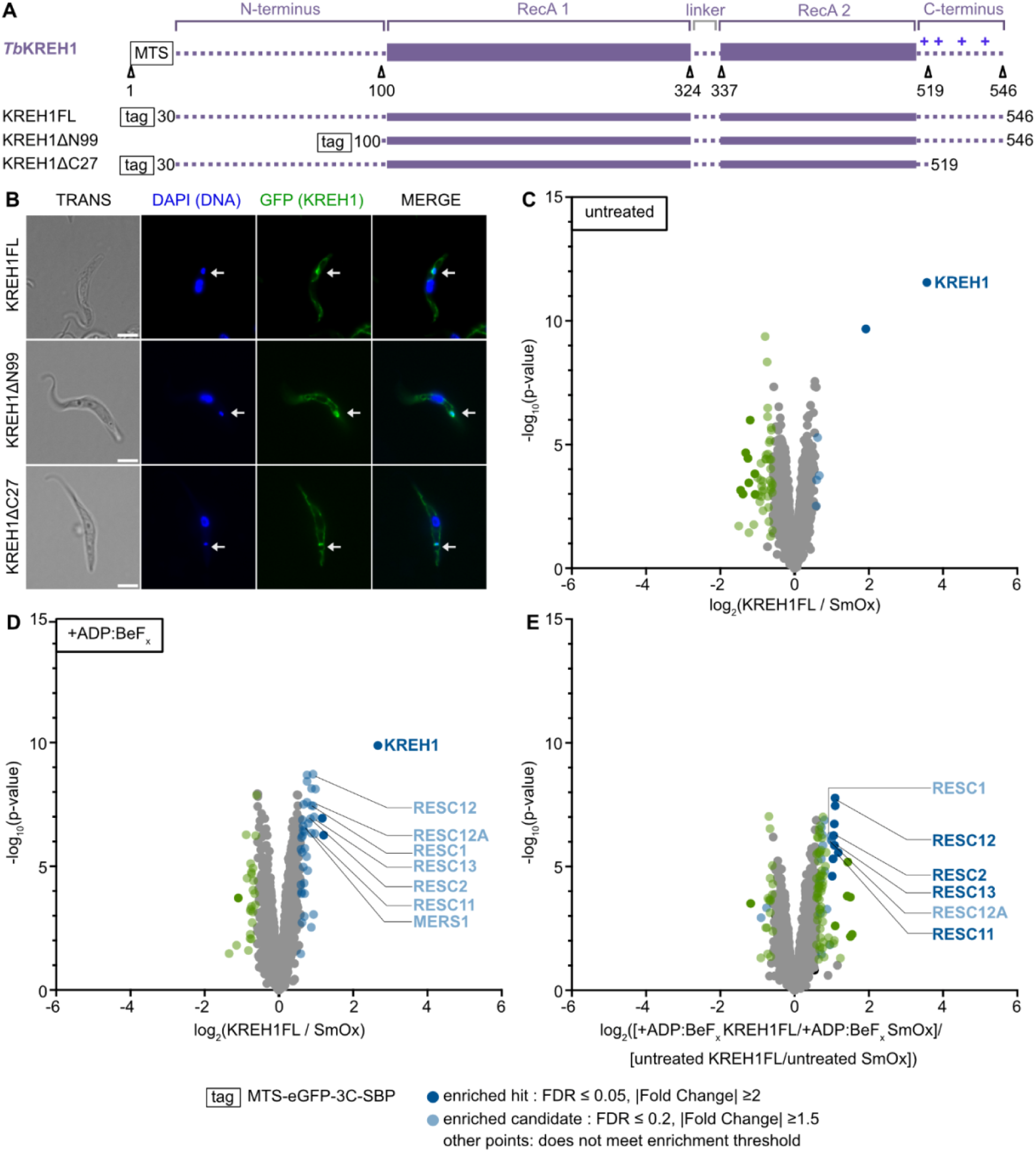
Differential quantitative mass spectrometry analysis of proteins co-purifying with tagged KREH1. **(A)** Design of KREH1 constructs with N-terminal purification tag (boxed) used for in vivo interactome analysis. Blue ‘+’ signs indicate patches in the sequence enriched with positive amino acids. (**B**) Subcellular localization of KREH1 24 h after induction. Fluorescence microscopy images of different KREH1 versions tagged with eGFP (green). Nuclei and kinetoplast (mitochondrial genome) are stained with DAPI (blue). The merged image shows the overlap of DAPI and eGFP signals. The corresponding transmitted light images are shown alongside. Scale bar = 5 µm. The white arrow indicates the kinetoplast. (**C**) Differential quantitative mass spectrometry analysis of tagged KREH1 compared to SmOx (Supplementary Table S5). (**D**) Differential quantitative mass spectrometry analysis of tagged KREH1 in the presence of the non-hydrolyzable ATP analog ADP:BeF_x_ compared to SmOx in the presence of ADP:BeF_x_ (Supplementary Table S6). (**E**) Differential analysis of (D) versus (C), i.e. the ADP:BeF_x_-dependent change in the KREH1FL interactome (Supplementary Table S7). Volcano plots display log_2_(fold-change) values calculated from TMT intensities using a moderated t-test (limma package, Ritchie et al. 2015). **Significance thresholds:** Proteins were annotated as hits at a false discovery rate (FDR) ≤ 0.05 and |fold change| ≥ 2, and as candidates at FDR ≤ 0.2 and |fold change| ≥ 1.5. Independently, proteins were annotated as enriched if they were a hit or candidate with a positive fold change in the corresponding comparison of the tagged construct against untagged SmOx. Colors: dark blue, enriched hit; light blue, enriched candidate; dark green, hit that is not enriched; light green, candidate that is not enriched; black, enriched but not significant in this comparison; gray, all remaining proteins. Note that the y-axis shows −log_10_(p-values) whereas the hit/candidate cut-offs are applied to the FDR. Enriched proteins of the RNA editing editosome complexes are specifically labeled.

We then hypothesized that additional RNA-mediated contacts might stabilize the interaction between KREH1 and the other proteins of the editing machinery. We aimed to trap KREH1 in an RNA-bound state by adding the adenosine triphosphate (ATP) analog adenosine diphosphate beryllium fluoride complex (ADP:BeF_x_) during cell lysis. ADP:BeF_x_ was chosen as it efficiently inhibited KREH1 dsRNA unwinding activity *in vitro* (Figure 5A). In the presence of ADP:BeF_x_, we were able to detect significant enrichment of components of the RESC complex such as RESC1, RESC2, RESC11, RESC12, RESC12A and RESC13 together with KREH1FL (Figure 2D, E and Supplementary Tables S6 and S7). In contrast, subunits of the catalytic RECC complex were not enriched, although technically detectable in the eluates (Supplementary Figure S3, Supplementary Table S8), indicating that KREH1 specifically associates with RESC modules. Additionally, our data indicated an interaction of KREH1 with MERS1, a subunit of the 5′ end pyrophosphohydrolase complex (PPsome), which binds mRNA 5′ ends (Sement et al. 2018; Li et al. 2011). The selective enrichment of the KREH1 interactome upon helicase inhibition further supports an RNA-dependent association with the editing machinery, consistent with previous findings (Hashimi et al. 2008; Panigrahi et al. 2003, 2006; Li et al. 2011).

### The KREH1 C-terminus mediates interaction with the RESC complex *in vivo*

Next, we sought to characterize the interactomes associated with N- and C-terminal regions of KREH1 through truncated protein variants, all in the presence of ADP:BeF_x_. The interactome of KREH1ΔN99 was comparable to that of the full-length construct, suggesting a limited role of the N-terminus in mediating protein-protein interactions under the experimental conditions used (Figure 3A; Supplementary Figure S4A and Supplementary Tables S9 and S10). In contrast, KREH1ΔC27 had reduced RESC co-purification, indicating that the C-terminal flexible region plays a role in the association of KREH1 with the mRNA editing machineries (Figure 3B, C; Supplementary Figure S4B and Supplementary Tables S11-13). The three constructs were recovered at different levels in the eluates (Supplementary Figure S2): relative to KREH1FL, KREH1ΔN99 was recovered ∼4.5-fold and KREH1ΔC27 ∼2.6-fold more abundantly. Importantly, KREH1ΔC27 was therefore present at a higher level than KREH1FL while co-purifying fewer RESC subunits, so the reduced RESC association of KREH1ΔC27 cannot be explained by a lower amount of bait in the pull-down. In conclusion, our KREH1 interactome analysis revealed associations between KREH1 and components of the RESC complex (Figure 3D, Supplementary Table S14); under our experimental conditions, these interactions are stabilized when KREH1 helicase activity is inhibited through a non-hydrolyzable ATP analog. Furthermore, our proteomics data indicate that the flexible C-terminus of KREH1 is an important molecular determinant mediating these interactions under RNA-engaged conditions.

**Figure 3.**
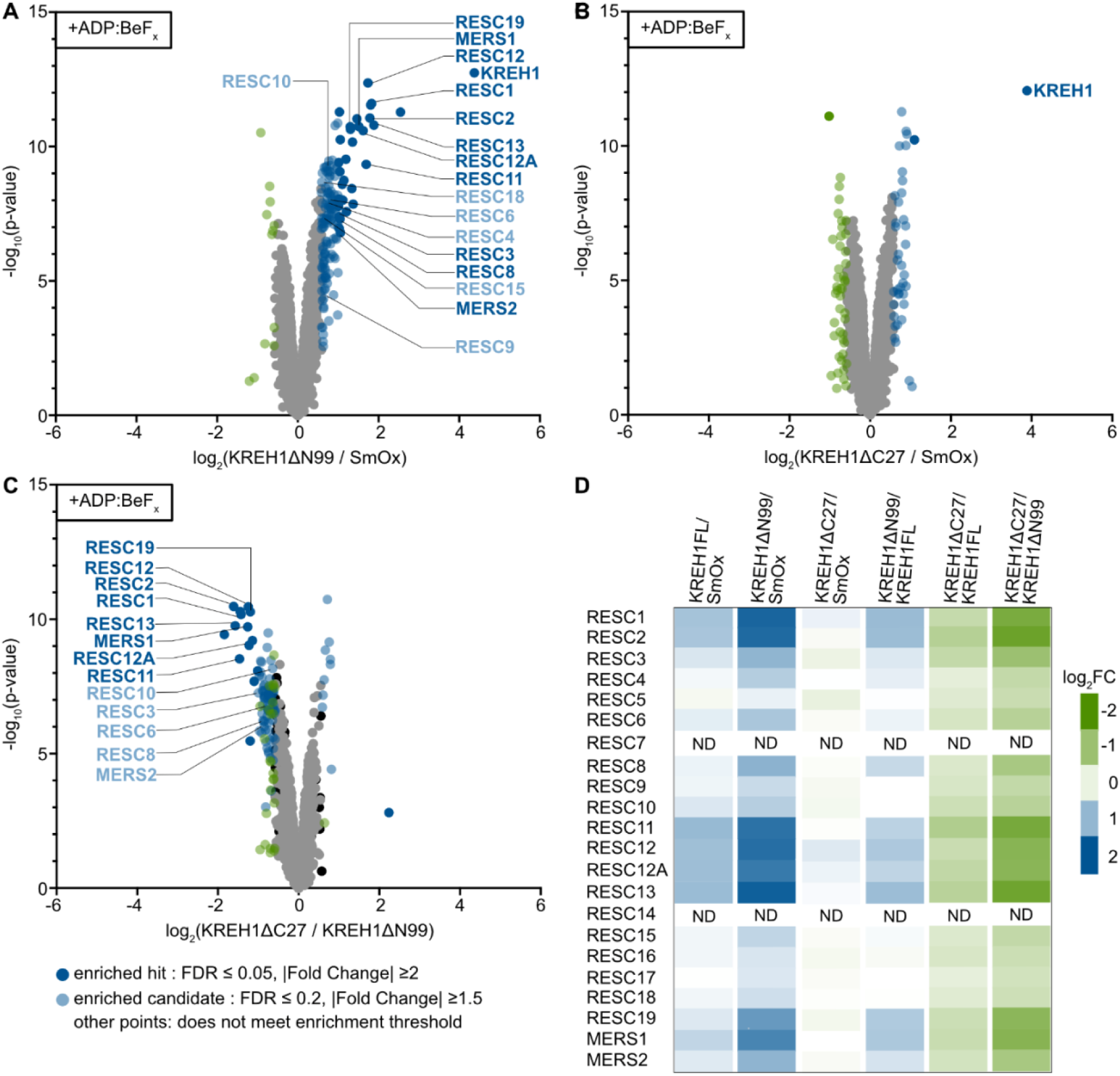
Differential quantitative mass spectrometry analysis of the proteins co-purifying in the presence and absence of the flexible N- and C-termini of KREH1. (**A**) KREH1ΔN99 compared to SmOx (Supplementary Table S9). (**B**) KREH1ΔC27 compared to SmOx (Supplementary Table S11). (**C**) KREH1ΔN99 compared to KREH1ΔC27, each normalized against SmOx (Supplementary Table S12). All samples and controls in the presence of ADP:BeF_x_. Volcano plots display log_2_(fold-change) values calculated from TMT intensities using a moderated t-test (limma package, Ritchie et al. 2015). **Significance thresholds**: Proteins were considered enriched hits (dark blue) at false discovery rate (FDR) ≤ 0.05 and fold change enrichment ≥ 2; enriched candidates (light blue) were defined at FDR ≤ 0.2 and fold change enrichment ≥ 1.5. Other proteins did not reach enrichment threshold. Enriched proteins of the RNA editing editosome complexes are specifically labeled. (**D**) Log_2_FC values for all RESC and PPsome subunits across different comparisons of KREH1 truncations against *T. brucei* SmOx and against each other (normalized against SmOx). Log_2_FC values are color-coded on a continuous scale from −2 (green) through 0 (white) to +2 (blue); the panel does not encode statistical significance (Supplementary Table S14). ND = subunit not detected in the eluates.

### The KREH1 C-terminus is important for RNA binding

Our co-proteome analysis revealed that the C-terminus of KREH1 is important for its *in vivo* association with the RNA editing machinery, thereby motivating a detailed characterization of the RNA-binding properties of KREH1 *in vitro*. For this, we expressed different recombinant KREH1 versions in *E. coli* and purified them (Figure 4A; Supplementary Figure S5A-M). Recombinant full-length KREH1 was insoluble, and deletion of at least 84 N-terminal residues was required to obtain a soluble protein. Even after truncation, the protein remained sensitive to NaCl concentration in the buffer and could not be concentrated beyond 3 mg/ml, thereby constraining the experimental conditions. We first assessed RNA-binding of KREH1ΔN99 to a synthetic mimic of the fully edited mRNA:gRNA duplex, based on the NADH dehydrogenase subunit 7 (ND7), an editing target affected by KREH1 knockout (Figure 4B) (Dubey et al. 2023). In a fluorescence polarization assay, we used an established ATPase-deficient helicase mutant in which the catalytic glutamate is substituted by glutamine (KREH1-E269Q); additionally, we supplemented the reaction with the ATP analog AMP-PNP to promote stable duplex binding without unwinding. The area under the curve (AUC) was calculated for each replicate as an alternative to determining a dissociation constant (*K_d_*), which was not feasible due to protein concentration limitations; the AUC was used to assess statistical differences in RNA binding. First, we focused on the N-terminus: further shortening of the N-terminal flexible region (KREH1ΔN84, KREH1ΔN94, KREH1ΔN99) did not affect the dsRNA binding capacity of KREH1 (Figure 4C, D), suggesting that these residues are not involved in RNA binding. Next, we tested variants with progressive C-terminal truncations: KREH1ΔN99ΔC12, KREH1ΔN99ΔC18 and KREH1ΔN99ΔC27, lacking 12, 18 or 27 C-terminal residues, respectively; these bound dsRNA with gradually lower affinity, indicating that the C-terminus contributes to RNA binding (Figure 4E, F; Supplementary Figure S6). For both N- and C-terminal truncations, single-stranded RNA (ssRNA) was bound only weakly. *In vitro* RNA-binding experiments with recombinant *T. brucei* KREH1 indicate that the flexible C-terminus of KREH1 is a molecular determinant that contributes to RNA binding.

**Figure 4.**
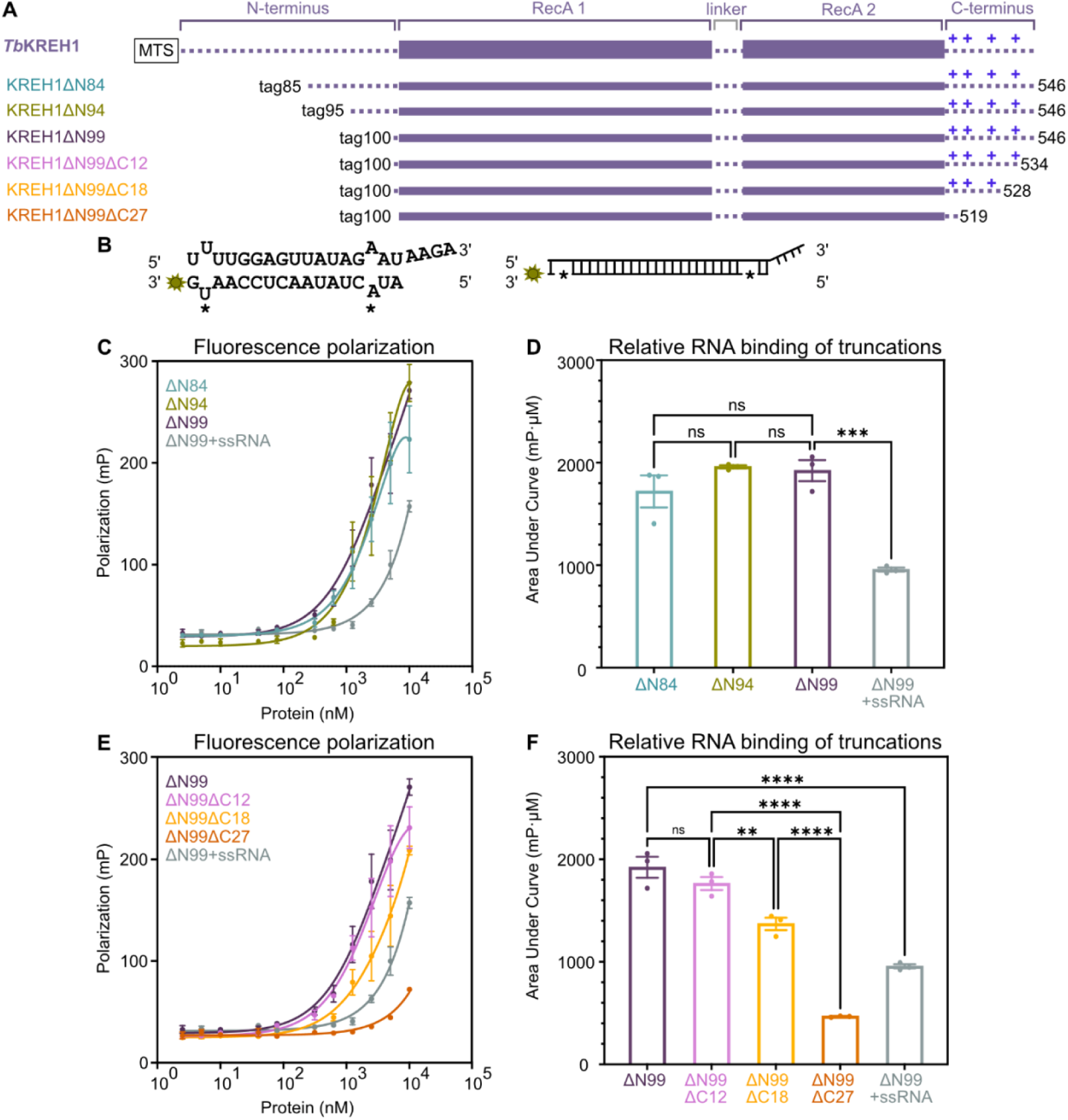
RNA-binding of KREH1 truncations. All constructs carry the E269Q ATPase-deficient mutation. **(A)** Schematic of KREH1 constructs used for biochemical characterization. (**B**) *Sequence of the double-stranded RNA (dsRNA) substrate derived from fully edited ND7 mRNA, containing two mismatches (**\****) and a 3′ overhang, and its schematic representation; the 3′ FAM label is indicated as a green star. (**C**) Fluorescence polarization curves of KREH1 N-terminal truncations. Data points from three technical replicates (n = 3) are plotted as mean ± standard deviation. (**D**) Statistical analysis of the Area Under the Curve (AUC) ± SEM from three technical replicates of (C) (n = 3). (**E**) Fluorescence polarization curves of KREH1 C-terminal truncations. Data points from three technical replicates (n = 3) are plotted as mean ± standard deviation. (**F**) Statistical analysis of the Area Under the Curve (AUC) ± SEM from three technical replicates of (E) (n = 3). Statistical significance between truncations was assessed by one-way ANOVA with Tukey’s multiple comparison test: ****p < 0.0001, ***p < 0.001, **p < 0.01, ns = not significant.

**Figure 5.**
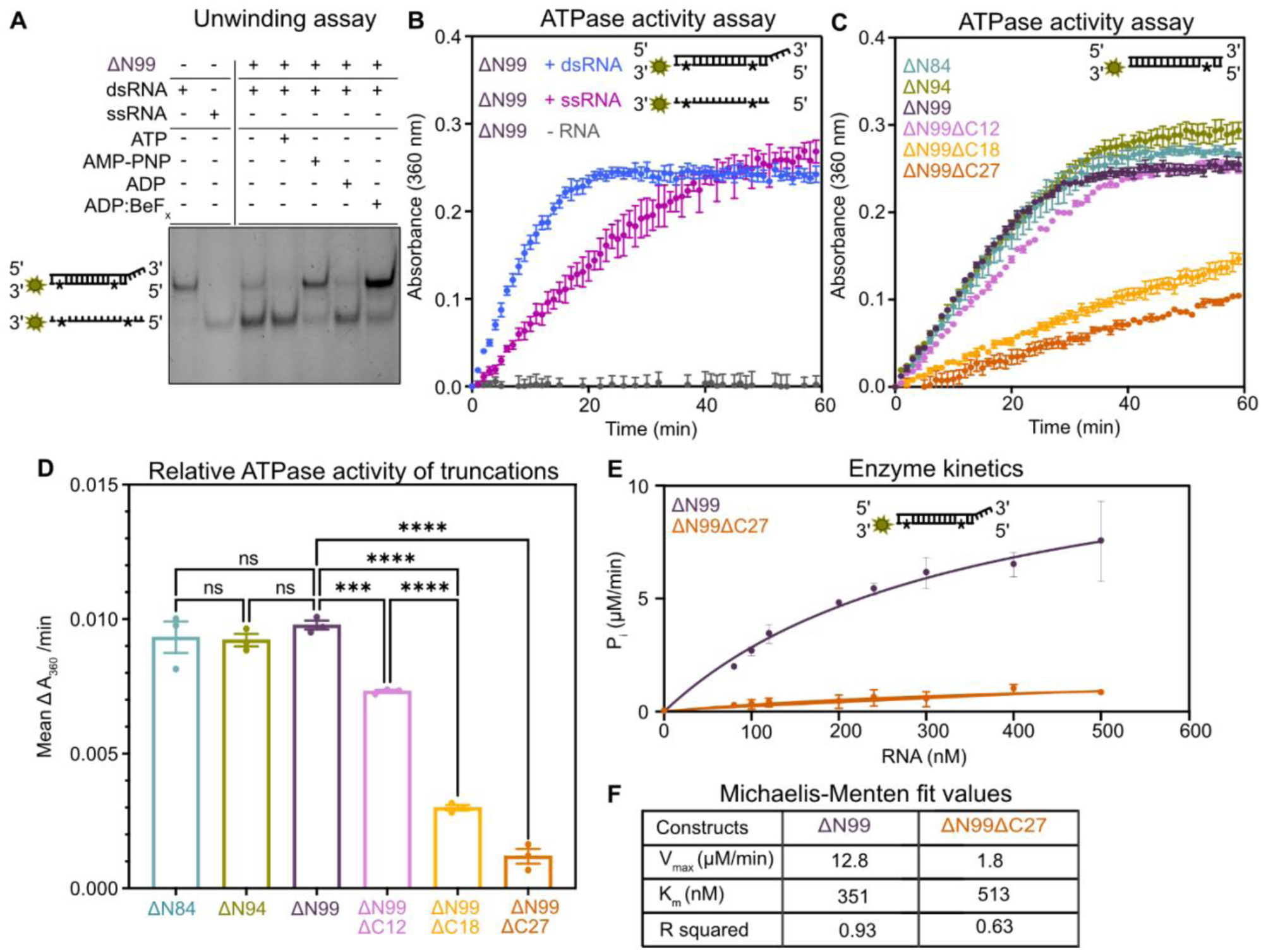
RNA unwinding and ATPase activity of KREH1. (**A**) RNA unwinding assay in the presence of different ATP analogs. One strand of the dsRNA substrate is FAM-labeled, the reaction is resolved by native PAGE, and detected with a fluorescein filter. (**B, C**) ATPase activity curves of KREH1ΔN99 in the presence of dsRNA (pink), ssRNA (black), or without RNA (gray), and ATPase activity curves of different KREH1 N- and C-terminal truncations, respectively. Data from three technical replicates (n = 3) are plotted as mean ± standard deviation. Schematic substrates are shown for each experiment; base mismatches are indicated by black stars; the green star indicates the 3′-FAM label. **(D)** Statistical analysis of (C). Bar plots of mean initial ATPase activity values, calculated for each replicate from the slope of the linear phase of ATPase curves (slope = Δ absorbance at 360 nm/min) (Supplementary Figure S7). Error bars indicate SEM (n = 3). Statistical differences in ATPase rates between truncations were assessed by one-way ANOVA with Tukey’s multiple comparison test: ****p < 0.0001, ***p < 0.001, ns = not significant. **(E)** Michaelis-Menten kinetics of dsRNA-induced ATPase activity of KREH1ΔN99 and KREH1ΔN99ΔC27. ATPase rates were measured at increasing dsRNA concentrations (nM) and are plotted as mean ± SD from three technical replicates (n = 3). Data were fitted by nonlinear regression in GraphPad Prism to obtain V_max_ and apparent K_m_ values. **(F)** Summary table of best-fit V_max_ and K_m_ values with R² from (E).

### *T. brucei* KREH1 is an RNA-dependent ATPase that unwinds dsRNA *in vitro*

In the absence of biochemical studies with purified *T. brucei* KREH1, we next evaluated its dsRNA unwinding activity using a strand-separation assay. One strand of the duplex substrate was labeled with a 3′ fluorophore, and unwinding was assessed qualitatively by comparing band shifts in a non-denaturing polyacrylamide gel electrophoresis (PAGE) mobility assay. In the presence of ATP, the reference construct KREH1ΔN99 fully separated the strands. When the reaction was supplemented with the non-hydrolyzable analogs AMP-PNP or ADP:BeF_x_, the RNA strand separation activity of KREH1 was inhibited, indicating that ATP-derived energy is necessary for RNA unwinding by *T. brucei* KREH1 (Figure 5A). Next, in an ATPase activity assay, the addition of dsRNA led to robust ATP turnover, whereas no activity was observed in the absence of RNA. Additionally, moderate ATPase activity was observed in the presence of ssRNA, consistent with the unwinding assay (Figure 5B). Our findings in *T. brucei* extend previous observations in *Leishmania major*, demonstrating that RNA-dependent ATPase activity is a conserved feature of KREH1 across kinetoplastids (Li et al. 2011).

Next, we performed ATPase and unwinding assays using our panel of KREH1 truncation constructs and a blunt-ended dsRNA substrate mimicking ND7 mRNA with a single mismatch. All N-terminal KREH1 truncations retaining the complete C-terminus exhibited ATPase activity comparable to that of the reference construct, KREH1ΔN99 (Figure 5C, D; Supplementary Figure S7). In contrast, the progressive shortening of the KREH1 C-terminus resulted in a gradual decrease in ATPase activity. The construct-dependent differences in ATP hydrolysis were mirrored by a similar pattern in RNA unwinding activity, suggesting that the C-terminus contributes to KREH1’s function (Supplementary Figure S8A-F). These findings are consistent with the RNA-binding assays, in which C-terminal truncation impaired binding. Further steady-state analysis of KREH1 ATP hydrolysis revealed that the reference construct KREH1ΔN99 turns over the substrate approximately sevenfold faster, with a V_max_ of 12.8 µM P_i_/min, compared to the construct with the shortest C-terminus (KREH1ΔN99ΔC27) exhibiting a V_max_ of 1.8 µM P_i_/min (Figure 5E, F). Collectively, these activity assays establish KREH1 as an RNA-dependent ATPase that unwinds RNA duplexes, with strand separation inhibited by non-hydrolyzable ATP analogs; the *in vitro* activity data further highlight the major contribution of the KREH1 C-terminus to function.

### KREH1 has a preference for dsRNA stems containing mismatches and 3′ overhangs

Next, we sought to identify specific RNA motifs with altered binding or activity of KREH1. To this end, we generated a series of derivatives of the previously used synthetic ND7 mRNA:gRNA duplex. We assessed the ATPase activity of KREH1ΔN84 on substrates with either perfect complementarity, defined mismatches, blunt or staggered ends, or combinations of these features; KREH1ΔN84 binds dsRNA indistinguishably from the reference construct KREH1ΔN99 and shows comparable patterns in the ATPase assay. Based on the activity curves, we categorized the substrate preferences into three groups: (i) fully base-paired RNA stems were the least effective substrates, regardless of whether overhangs were present; (ii) substrates with mismatches were unwound with intermediate efficiency; (iii) substrates carrying a 3′ overhang with a mismatch in direct proximity were processed with the highest efficiency (Figure 6A, B; Supplementary Figure S9A, B). When expanding our substrate pool beyond pure RNA duplexes, we found that KREH1 was also moderately active on RNA:DNA hybrids; similarly, mismatches promoted processing in this context. Pure dsDNA duplexes did not function as substrates regardless of mismatches, suggesting a preference for A-form helix substrates (Figure 6C, D). In conclusion, our analysis of substrate preferences reveals that KREH1 can act on diverse dsRNA substrates and even DNA:RNA hybrids; however, KREH1 exhibits the highest efficiency on dsRNA with mismatches near 3′ overhangs.

**Figure 6.**
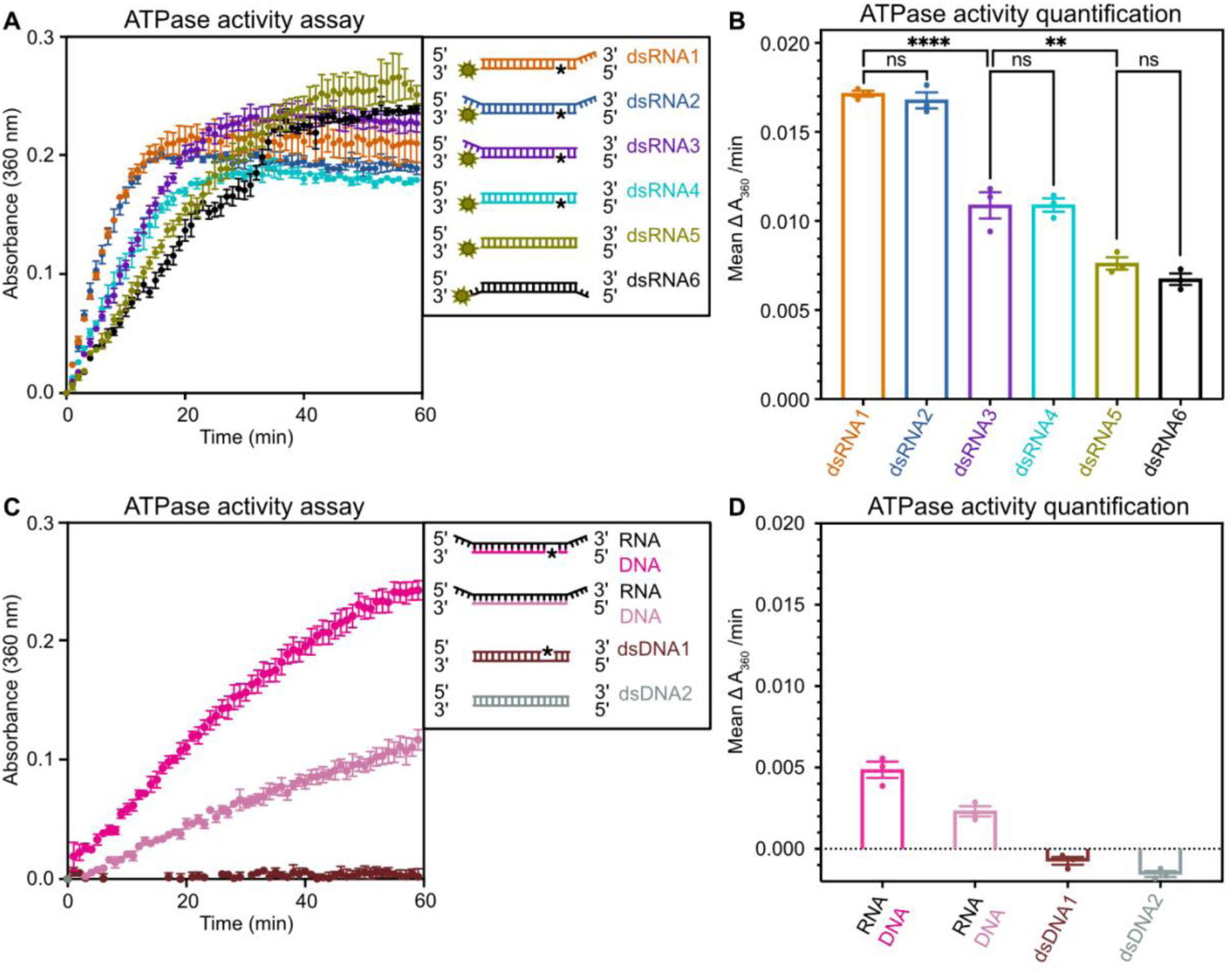
Substrate preferences of KREH1. **(A)** ATPase activity of KREH1ΔN85 on different dsRNA substrates. Linear schematics of the oligos, color coded to match the curves, indicate structural features such as overhangs and mismatched base pairs (*); three technical replicates (n = 3) are plotted as individual data points ± standard deviation from mean. (**B**) Comparison of the ATPase activities from (A): Bar plots show mean initial ATPase activity values ± SEM from three technical replicates (n = 3). Initial rates were calculated for each replicate from the slope of the linear phase of the ATPase curve (slope = Δ absorbance at 360 nm/min) (Supplementary Figure S9A). Statistical significance was assessed by one-way ANOVA with Tukey’s multiple comparison test: ****p < 0.0001, **p < 0.01, ns = not significant. (**C**) ATPase activity of KREH1ΔN85 with RNA:DNA hybrid and dsDNA oligos; three technical replicates (n = 3) are plotted as individual data points ± standard deviation from mean. Linear schematics of the oligos, color coded to match the curves, indicate structural features such as overhangs and mismatched base pairs (*). **(D)** Quantification of ATPase activity from (C): Bar plots of mean initial ATPase activity values, calculated for each replicate from the slope of the linear phase of ATPase curves (slope = Δ absorbance at 360 nm/min). Error bars indicate SEM (n = 3) for (C) (Supplementary Figure S9B).

## Discussion

RNA editing associated helicases drive remodeling and progression of the kinetoplastid mitochondrial mRNA editing machinery via ATP hydrolysis. A detailed understanding of the molecular functions of the individual KREH helicases is necessary to elucidate the mechanisms underlying pan-editing, the remarkable process that recodes extensive regions of mRNA. Our study focused on the DEAD-box helicase KREH1 and identified a key role for its flexible C-terminal tail in association with the RESC complex, as removal of the tail weakened the KREH1 interactome in cells. A similar mechanism had been observed for the mitochondrial helicase CYT19 from *Neurospora crassa*, for which mutations in the C-terminal tail altered RNA binding and ATPase activity (Busa et al. 2017). *In vitro* RNA-binding assays and ATPase activity measurements further indicated that the C-terminus is an important molecular determinant contributing to these activities and that inhibition of ATPase activity stalls KREH1 on RNA. Together, the *in vitro* and *in vivo* data suggest that the interaction between KREH1 and the editing machinery is at least partly dependent on the RNA-bound state of KREH1. Our results are consistent with prior studies proposing an RNA-dependent interaction between KREH1 and the RESC or RECC complexes (Li et al. 2011). Within the KREH1 C-terminus, a conserved cluster of positively charged residues can provide affinity towards the negatively charged phosphates of the RNA backbone. Compared with the *T. brucei* enzyme analyzed here, the C-terminal tail in *Leishmania* species is longer but retains a similar positive charge distribution, suggesting a conserved function (Figure 1C). In light of the importance of the KREH1 C-terminal tail, prior results obtained with KREH1 versions modified with C-terminal tags may warrant nuanced re-evaluation.

In contrast to a previous study reporting associations with both RESC and RECC (Dubey et al. 2023), our KREH1 interactome analysis predominantly identified RESC subunits, with little to no association with RECC components; however, RNA editing complexes are highly heterogeneous, and associations may be highly context-specific. Within the RESC complexes, enrichment was slightly higher across all samples for the RESC1-RESC2 heterodimer, as well as for the RESC11, RESC12/12A, RESC13, and RESC19 subunits (Figure 3D). When placing these results into a topological context, a previously proposed model of the RESC complex bound to pre-edited mRNA and a gRNA:mRNA duplex (termed RESC-B) provides a useful framework (Liu et al. 2023). This model, based on single-particle cryo-electron microscopy, proteomics, and enhanced *in vivo* UV cross-linking/affinity purification/RNA sequencing (eCLAP) data, suggests that the 5′ region of the pre-edited mRNA enters RESC-B through peripheral subunits, including RESC11, RESC12/12A, RESC13, and RESC9, whereas the 3′ region of the edited mRNA, engaged in a duplex with a gRNA, exits through RESC5 and RESC6 (Liu et al. 2023). Our proteomics data would place KREH1 near the 5′ end of the pre-edited mRNA entry site in RESC-B. This localization is consistent with a role in early stages of RNA editing initiation but contrasts with the hypothesis that KREH1 may directly engage gRNA:mRNA duplexes, which are predicted to reside on the opposite side of the RESC-B structure. The RESC1-RESC2 heterodimer has also been identified as part of a ‘storage’ particle stabilizing gRNA hairpins, termed RESC-A, together with RESC3, RESC4, RESC5, and RESC6 (Liu et al. 2023). As the other subunits contained in this particle are not enriched in our dataset, and given our observed substrate preferences, we find no evidence that KREH1 would specifically function within the RESC-A context. The selective enrichment of RESC1-RESC2 in the KREH1 interactome may instead suggest an interaction context in which the RESC1-RESC2 heterodimer operates independently of the rest of the RESC complex (Dolce et al. 2023; Dubey et al. 2021; Liu et al. 2023; Madina et al. 2015). An interaction network of the mRNA 5′-end processing ‘PPsome’, which includes the NUDIX hydrolase MERS1, MERS2, as well as RESC19 (formerly MERS3), RESC1, RESC2, RESC12A (formerly REMC5A), and RESC13 (formerly TbRGG2) was proposed earlier (Sement et al. 2018). Components of this previously reported network appear to be partially represented in the KREH1 interactome, potentially pointing towards a context-specific RESC-associated particle or alternative, currently poorly characterized, RESC configurations, a possibility that warrants further investigation. Overall, the dynamic and heterogeneous nature of RNA editing complexes likely complicates the spatial interpretation of interactome data. Resolving these discrepancies will require higher-resolution approaches, such as cross-linking mass spectrometry, single-particle cryo-electron microscopy, or *in situ* structural methods, to validate direct protein-protein interactions.

From the perspective of substrate preference, our *in vitro* analyses indicate that KREH1 exhibits a broad substrate range, with the highest ATPase activity observed for imperfect RNA duplexes containing 3′ overhangs and nearby mismatches. We speculate that the proposed RESC-A ‘gRNA storage’ particle would be unlikely to present a suitable substrate for KREH1, as within RESC-A, both gRNA ends appear occluded: the 5′ triphosphate is capped by RESC2 (Dolce et al. 2023; Liu et al. 2023), and the 3′ end is shielded by RESC5 and RESC6 (Liu et al. 2023), consistent with the underrepresentation of the corresponding RESC subunits in our KREH1 interactome data. In contrast, within the larger RESC particle engaged with the mRNA:gRNA duplex (RESC-B), the 3′ mRNA end next to the anchor region remains accessible and could be targeted by KREH1. In this context, KREH1 may help fine-tune local RNA remodeling to favor the pairing of the cognate gRNA. We speculate that a functional connection or partial overlap with the role of RESC13 (TbRGG2), which is also enriched in our KREH1 interactome, is present: RESC13 has been shown to promote editing progression and modulate RNA-RNA interactions (Ammerman et al. 2010; Fisk et al. 2008; Simpson et al. 2017). Future analysis of the co-dependence and/or distinct functions of KREH1 with RESC11, RESC12, RESC12A and/or RESC13 will be necessary.

We conclude that *T. brucei* KREH1 is a RESC-associated, ATP-dependent RNA helicase capable of targeting a broad range of dsRNA substrates. Because KREH1-mediated RNA unwinding can be inhibited by ATP analogs, a measure that also enriches RESC subunits in an RNA-dependent manner, such strategic modulation may provide a valuable route for targeted purification of RESC particles and in-cell structural studies. Ultimately, these approaches could help to fully elucidate the mechanisms and dynamics of kinetoplastid pre-mRNA editing.

### Limitations of this study

**Technical limitations and open possibilities.** For the interactome analysis, full-length and truncated KREH1 constructs were overexpressed in non-clonal cell pools with large tags integrated into the rRNA locus. Additionally, the interactome analysis was performed using technical triplicates derived from the same cell lysate; therefore, the reported FDRs reflect technical rather than biological reproducibility. For the biochemical assays, the removal of N-terminal residues of KREH1 was necessary to obtain sufficient soluble protein. Therefore, the molecular function of the flexible N-terminus of KREH1 remains unaddressed and we cannot exclude the involvement of N-terminal residues in RNA binding and protein-protein interactions. Purified KREH1 required careful buffer optimization to maintain both protein stability and RNA-binding capacity, resulting in binding curves that did not reach saturation and thereby limiting absolute affinity measurements. For the same reason, the steady-state ATPase kinetics were acquired over a substrate range that does not reach saturation (up to 500 nM, against apparent K_m_ values of 350–510 nM) and at an enzyme concentration that is not negligible relative to K_m_; the reported V_max_ and K_m_ are therefore apparent values under these conditions, and the fit for KREH1ΔN99ΔC27 in particular is poorly constrained (R² = 0.63). PAGE-based assays allow only qualitative assessment of RNA unwinding activity. **Suggestions for further research.** Cross-interactome analysis of the identified interactors and interactome analysis of C-terminally tagged KREH1 constructs could be performed. Quantitative Förster resonance energy transfer (FRET)-based assays would provide further quantification of strand-separation activity.

### Data Availability

The mass spectrometry proteomics data have been deposited to the ProteomeXchange Consortium via the PRIDE partner repository with the dataset identifier PXD067570.

### Supplementary data statement

A supplementary folder with files containing Supplementary Figures and Supplementary Tables has been provided.

## Acknowledgements

We acknowledge Martin Pelosse for support in using the Eukaryotic Expression Facility at EMBL Grenoble. This work used the platforms of the Grenoble Instruct-ERIC center (ISBG; UAR 3518 CNRS-CEA-UGA-EMBL) within the Grenoble Partnership for Structural Biology (PSB), supported by FRISBI (ANR-10-INBS-0005-02) and GRAL, financed within the University Grenoble Alpes graduate school (Écoles Universitaires de Recherche) CBH-EUR-GS (ANR-17-EURE-0003). We thank Jennifer Schwarz from the EMBL Proteomics Core Facility for her support with the mass spectrometry data acquisition and analysis. We thank Sarah Kaspar and Eva Geissen in the Data Science Centre at EMBL for their support with biostatistics. We thank Mark Carrington and Suzanne McDermott for critical comments on the manuscript. The project was supported by a grant from the French Agence Nationale de la Recherche to E.K. (ANR-20-CE11-0016). The lab of Eva Kowalinski is supported by ERC grant TRANSPLIC, 101170068, funded by the European Union. Views and opinions expressed are, however, those of the authors only and do not necessarily reflect those of the European Union. Neither the European Union nor the granting authority can be held responsible for them. The authors thank the Kowalinski lab members for discussions, comments and constructive criticism throughout the project.

## Authors contributions

EK designed and supervised the study and acquired funding. RY and LT carried out experimental work. FS analyzed mass spectrometry data. RY and EK interpreted the data and wrote the manuscript.

