## Supplementary Data File for "The C-terminus of the KREH1 helicase is important for RNA binding and association with mitochondrial RNA editing complexes in *T. brucei*"

### Supplementary Figures

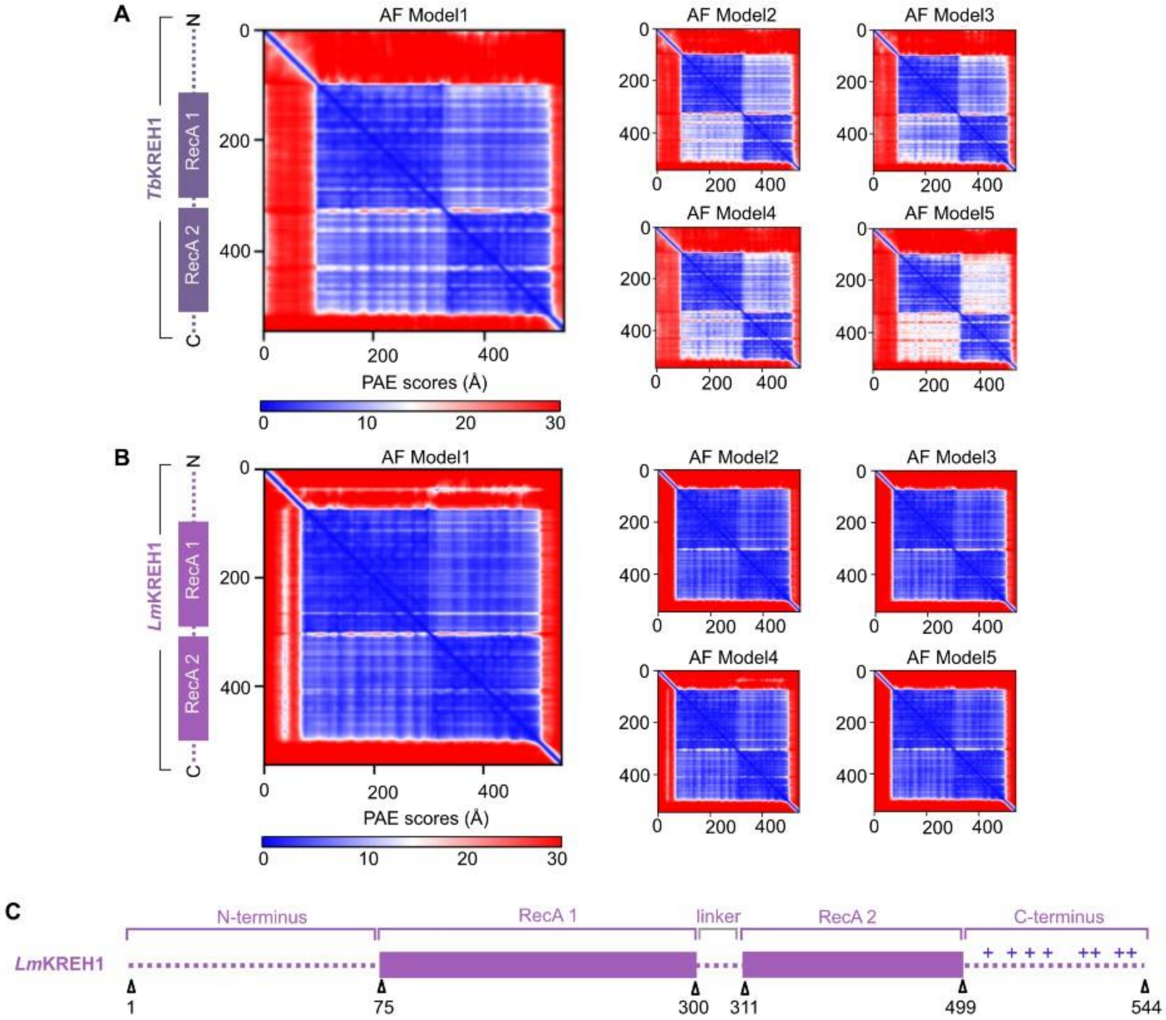


**Supplementary Figure S1. KREH1 topology across species.** (**A**) Predicted Alignment Error (PAE) plot of five AlphaFold models of KREH1 from Trypanosoma brucei (Tb) and (**B**) Leishmania major (Lm), showing pairwise positional uncertainty (blue: low, red: high, in Å). (**C**) Annotated linear schematic of full-length LmKREH1: a flexible amino-terminus, the helicase core consisting two RecA-like domains, and a flexible carboxy-terminus with positively charged residues (blue patches of plus sign).

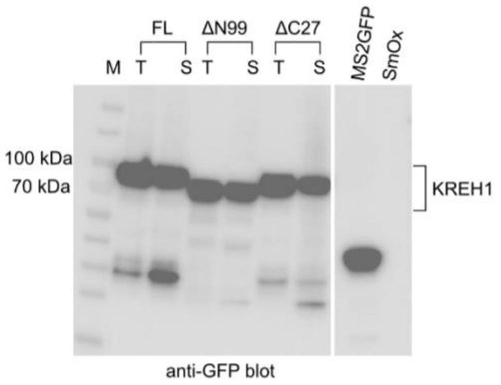


**Supplementary Figure S2. Western blot analysis of over-expressed KREH1.** Western blot against the eGFP present in total (T) and clarified soluble (S) lysates from cell lines expressing N-terminally MTS-eGFP-3C-SBP-fused KREH1FL, KREH1ΔN99 and KREH1ΔC27 KREH1 variants with T and S fractions showed side by side for each variant. MS2-GFP served as a positive control, and SmOx total cell lysate as a negative control on the same blot.

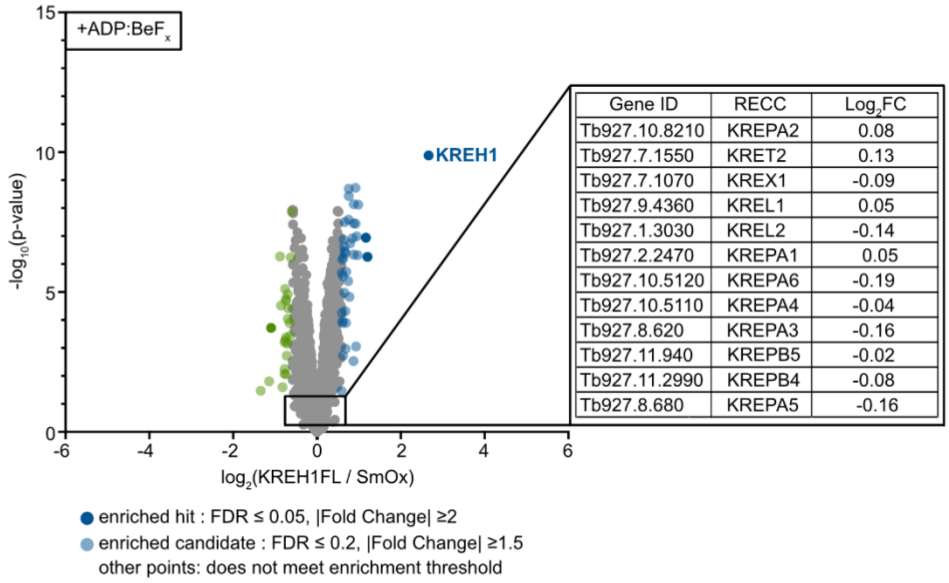


| Gene ID | RECC | $Log_2FC$ |
|---|---|---|
| Tb927.10.8210 | KREPA2 | 0.08 |
| Tb927.7.1550 | KRET2 | 0.13 |
| Tb927.7.1070 | KREX1 | -0.09 |
| Tb927.9.4360 | KREL1 | 0.05 |
| Tb927.1.3030 | KREL2 | -0.14 |
| Tb927.2.2470 | KREPA1 | 0.05 |
| Tb927.10.5120 | KREPA6 | -0.19 |
| Tb927.10.5110 | KREPA4 | -0.04 |
| Tb927.8.620 | KREPA3 | -0.16 |
| Tb927.11.940 | KREPB5 | -0.02 |
| Tb927.11.2990 | KREPB4 | -0.08 |
| Tb927.8.680 | KREPA5 | -0.16 |

**Supplementary Figure S3. Differential $log_2$(fold-change) analysis of RECC subunits.** Differential quantitative mass spectrometry analysis of tagged KREH1 compared to untagged SmOx, both samples in the presence of ADP:BeF$_x$. The box indicates that RECC subunits are in principle present and detectable, but not enriched with KREH1. All twelve RECC subunits listed were quantified in all 18 TMT channels with 2–7 razor peptides, and none reached the enrichment threshold. Volcano plots display $log_2$(fold-change) values calculated from TMT intensities using a moderated t-test (limma package, Ritchie et al. 2015). **Significance thresholds:** Proteins were considered enriched hits (dark blue) at false discovery rate (FDR) ≤ 0.05 and |Fold Change| ≥ 2; enriched candidates (light blue) were defined at FDR ≤ 0.2 and |Fold Change| ≥ 1.5. Other proteins did not reach enrichment threshold. Non-enriched proteins from RNA Editing Catalytic Complex are indicated in the table with $log_2$FC (inset) (Table S8).

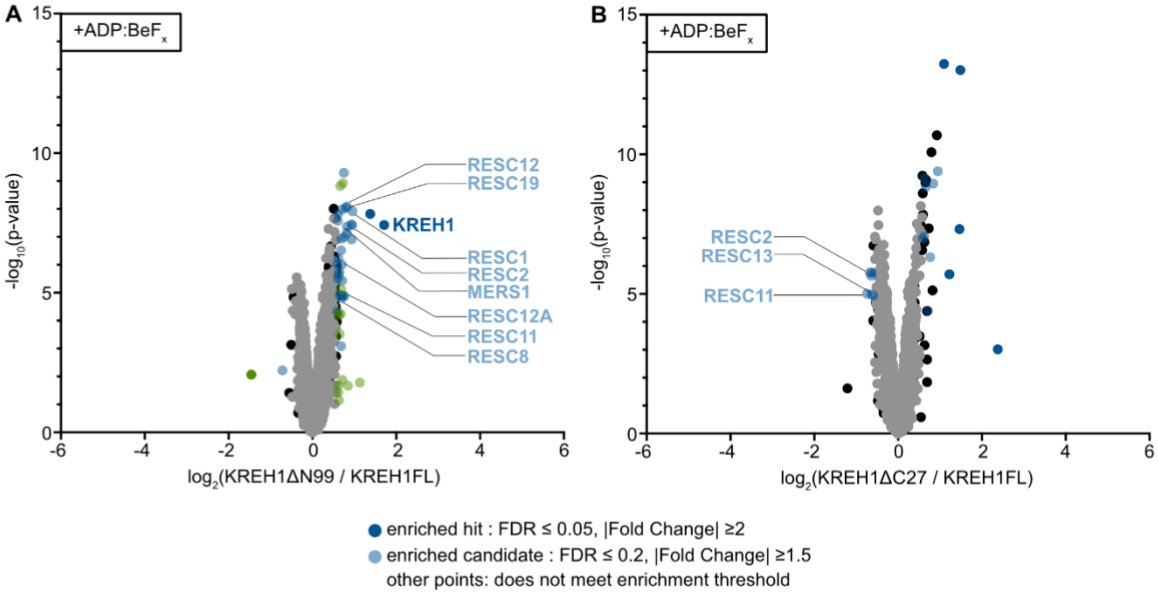


**Supplementary Figure S4. Differential quantitative mass spectrometry analysis of the proteins co-purifying with KREH1 N- and C- terminal truncations.** (**A**) KREH1ΔN99 (normalized against SmOx) compared to KREH1FL (normalized against SmOx) (Table S10). (**B**) KREH1ΔC27 (normalized against SmOx) compared to KREH1FL (normalized against SmOx) (Table S13). All samples and controls in the presence of ADP:BeF$_x$. Volcano plots display $\log_2$(fold-change) values calculated from TMT intensities using a moderated t-test (limma package, Ritchie et al. 2015). **Significance thresholds:** Proteins were considered enriched hits (dark blue) at false discovery rate (FDR) $\leq 0.05$ and |Fold Change| $\geq 2$; enriched candidates (light blue) were defined at FDR $\leq 0.2$ and |Fold Change| $\geq 1.5$. Other proteins did not reach enrichment threshold. Enriched proteins of the RNA editing editosome complexes are specifically labeled.

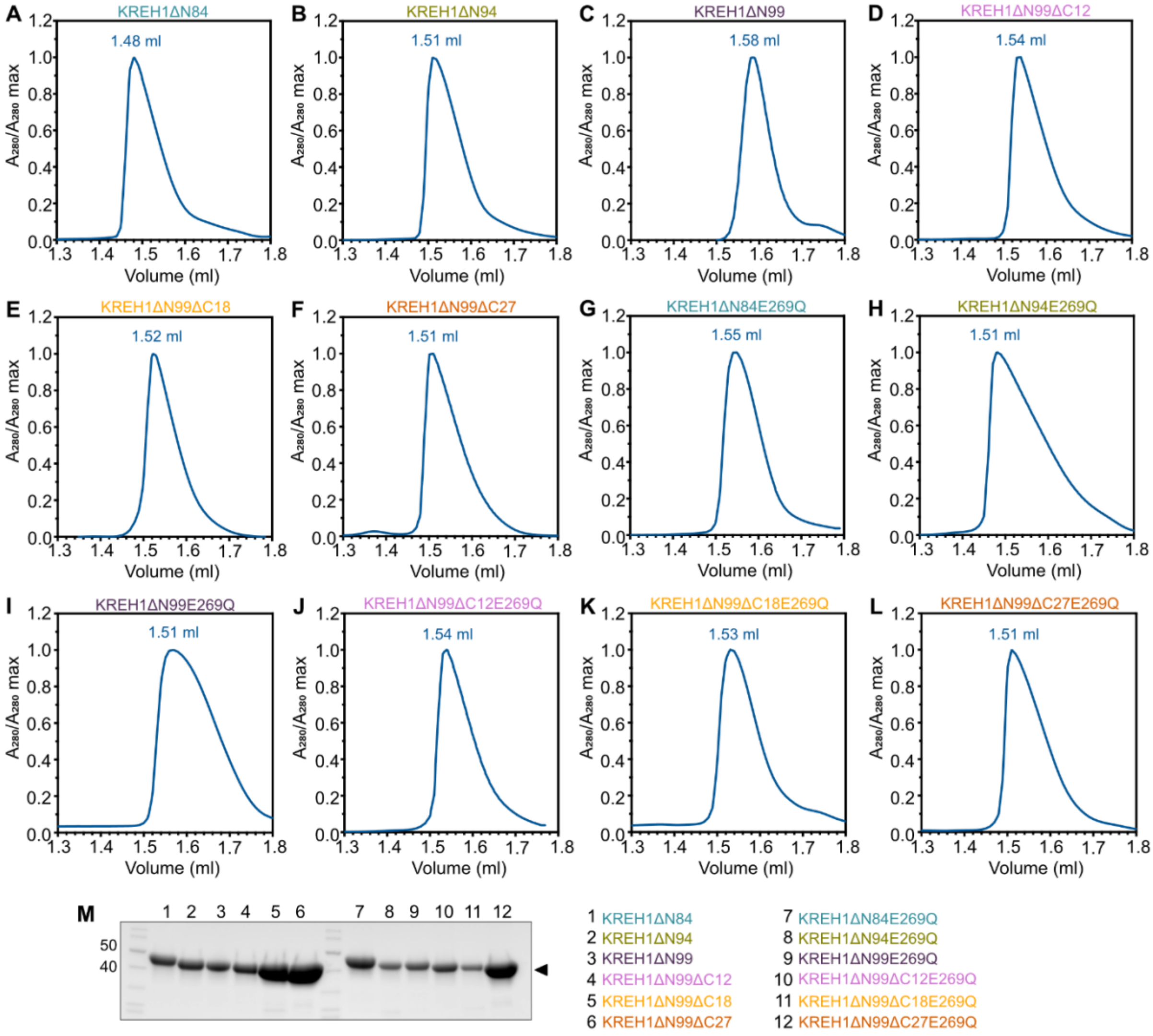


**Supplementary Figure S5. Purification of KREH1 constructs.** (**A-L**) SEC profiles of KREH1 on a Superdex 200 Increase 3.2/300 column, monitored at 280 nm (blue trace) and normalized to the peak maximum. The main peak corresponds to the homogeneous protein population, with retention volumes indicated above each peak. **(M)** Peak fractions were analyzed by 12% SDS-PAGE and visualized with Coomassie Blue staining.

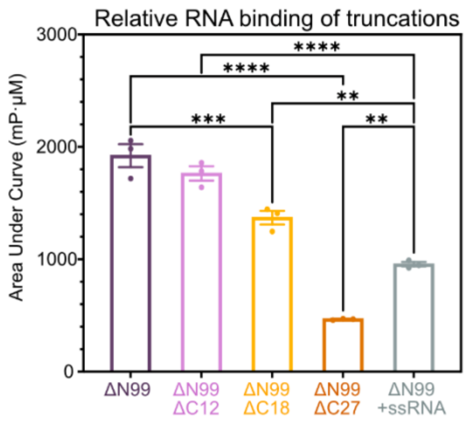


**Supplementary Figure S6. Additional statistical analysis of RNA-binding of KREH1 truncations.** Statistical analysis of the Area Under the Curve (AUC) ± SEM from three technical replicates from fluorescence polarization experiment of KREH1 C-terminal truncations (n = 3). Statistical significance between truncations was assessed by one-way ANOVA with Tukey's multiple comparison test: ****p < 0.0001, ***p < 0.001, **p < 0.01.

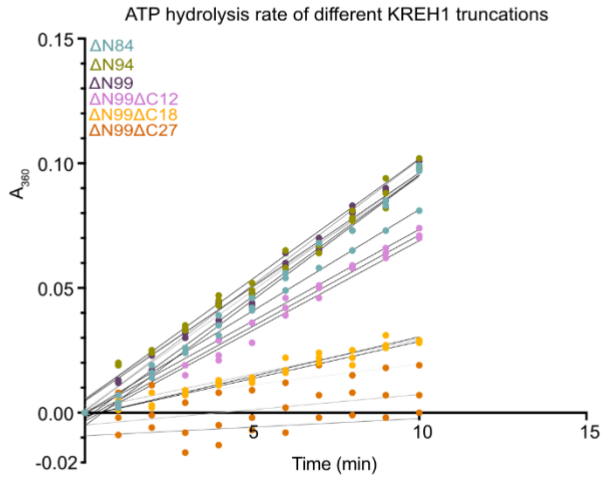


**Supplementary Figure S7. Time course of steady-state ATPase activity at constant substrate concentration for KREH1 truncations.** The solid line represents linear regression of the initial phase (0–10 min) to determine the initial ATPase rate (slope = Δ absorbance at 360 nm/min) of different KREH1 truncations in the presence of the same double-stranded RNA oligo substrate.

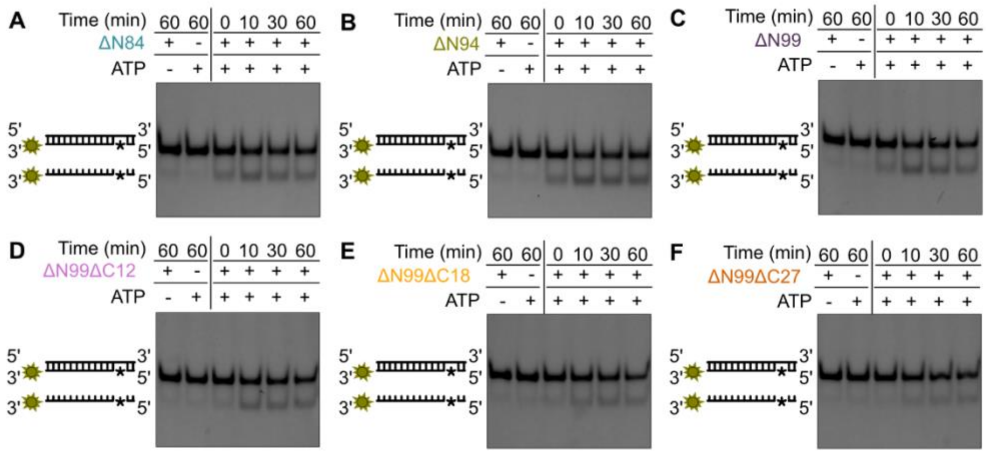


**Supplementary Figure S8. RNA unwinding analysis of KREH1 N- and C-terminal truncations. (A–F)** Unwinding assays of different KREH1 truncations using a FAM-labeled (green star) blunt-ended dsRNA substrate containing a single mismatched base pair (*). Reaction products were resolved on 20% native-PAGE and visualized with fluorescence imaging with a fluorescein filter.

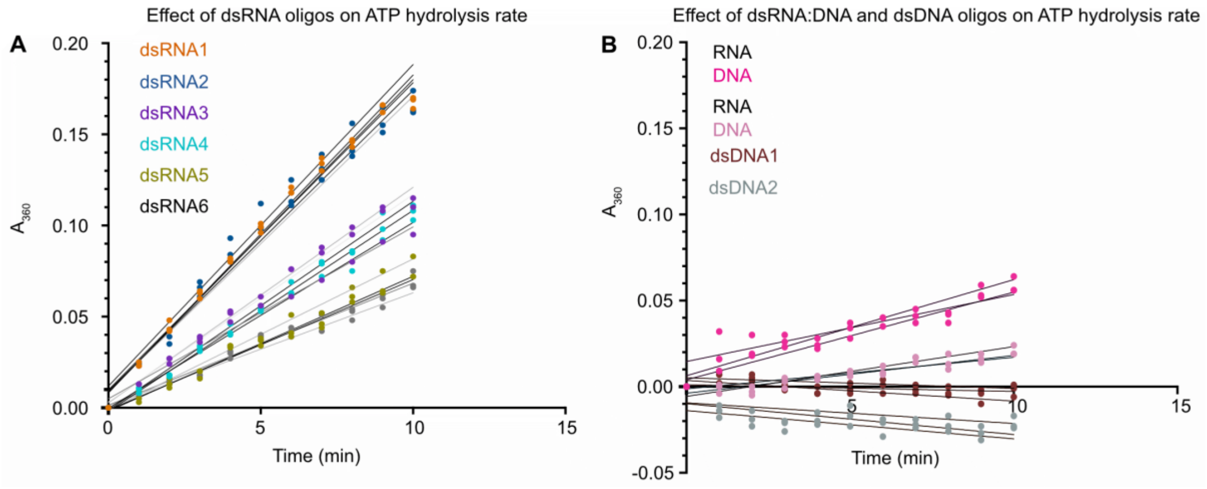


**Supplementary Figure S9. Time course of steady-state ATPase activity at constant substrate concentration for different substrates.** The solid line represents linear regression of the initial phase (0–10 min) to determine the initial ATPase rate (slope = Δ absorbance at 360 nm/min) for KREH1ΔN84 against (**A**) double-stranded RNA and (**B**) RNA:DNA hybrids and double-stranded DNA oligo substrates.

### Supplementary Tables

**The following titles correspond to the supplementary tables provided in the Supplementary_Tables.xlsx file.**

**Table S1** Details on plasmids used and clones generated in this study

**Table S2** Primers used for cloning and *Tb* cell line generation in this study

**Table S3** *Trypanosoma brucei* cell lines generated for this study

**Table S4** Sequences of oligonucleotides used in biochemical assays

**Table S5** Raw data values for $log_2$(FC) and -$log_{10}$(p-value) plotted in Figure 2C

**Table S6** Raw data values for $log_2$(FC) and -$log_{10}$(p-value) plotted in Figure 2D

**Table S7** Raw data values for $log_2$(FC) and -$log_{10}$(p-value) plotted in Figure 2E

**Table S8** Raw data values for $log_2$(FC) and -$log_{10}$(p-value) of the non-enriched RECC subunits shown in the inset table of Supplementary Figure S3

**Table S9** Raw data values for $log_2$(FC) and -$log_{10}$(p-value) plotted in Figure 3A

**Table S10** Raw data values for $log_2$(FC) and -$log_{10}$(p-value) plotted in Supplementary Figure S4A

**Table S11** Raw data values for $log_2$(FC) and -$log_{10}$(p-value) plotted in Figure 3B

**Table S12** Raw data values for $log_2$(FC) and -$log_{10}$(p-value) plotted in Figure

**Table S13** Raw data values for $log_2$(FC) and -$log_{10}$(p-value) plotted in Supplementary Figure S4B

**Table S14** Raw data values for $log_2$(FC) for RESC and PPsome complex subunits detected across KREH1 truncations plotted in Figure 3D

### Source Data

Source data for Figures 2B, 4C-F, 5A-F, 6A-D and Supplementary Figures S2, S5, S6, S7, S8 and S9 are provided in the Source_Data.xlsx file.
